# Restoring compromised Blood-Retina-Barrier integrity with Netrin-1 overexpression

**DOI:** 10.1101/2025.06.14.658755

**Authors:** Jessica Furtado, Thomas E. Zapadka, Hyojin Park, Kevin Boyé, Jonathan B. Demb, Anne Eichmann

## Abstract

The blood-retina barrier (BRB) protects retinal neuronal function and enables vision. A compromised, leaky BRB is a hallmark of vision-threatening retinal diseases such as diabetic retinopathy (DR) and wet age-related macular degeneration (AMD) that affect millions of persons worldwide. Strategies to enhance BRB integrity hold promise as therapeutic interventions to prevent vision loss. Previous studies identified Netrin-1 (*NTN1*) as a key regulator of BRB stability and revealed reduced Netrin-1 signaling in DR patients, suggesting that Netrin-1 supplementation could help preserve BRB function and prevent disease progression. Herein, we used genetic *NTN1* overexpression to investigate effects on BRB development and maintenance. We show that global *NTN1* overexpression converted leaky vessels at the P5 angiogenic front into a non-leaky state. In pathological settings, *NTN1* overexpression reinforced BRB integrity in oxygen-induced retinopathy (OIR), improving electroretinogram (ERG) amplitudes and rescued vascular leak in laser-induced choroidal neovascularization (CNV). Mechanistically, we determined that *NTN1* overexpression minimally and transiently affected retinal angiogenesis in an Unc5b independent manner, while Netrin’s barrier stabilizing effects were Unc5b dependent. These findings establish Netrin-1 as a promising therapeutic target for preventing BRB breakdown in retinal vascular diseases and suggest that reinforcing the Netrin-1/Unc5b signaling pathway may provide a strategy to selectively stabilize the BRB.

## INTRODUCTION

The blood-retina barrier (BRB) is a highly selective vascular barrier that tightly regulates the movement of molecules between the bloodstream and the neural retina, thereby maintaining the specialized microenvironment necessary for retinal function^1^. The BRB consists of an inner barrier formed by retinal endothelial cells (ECs) and an outer barrier formed by retinal pigment epithelial (RPE) cells, both of which utilize tight junctions to restrict permeability while allowing controlled nutrient and waste exchange. Disruption of the integrity of the inner BRB accelerates the development of multiple diseases including diabetic retinopathy (DR) and wet age-related macular degeneration (AMD) by leading to vascular leak, fluid accumulation and inflammation that cause neuronal damage and vision loss^2^. Similar breakdown of the blood-brain barrier is involved in diseases such as multiple sclerosis, Alzheimer’s disease and stroke^2–6^.

DR is the leading cause of vision loss worldwide, where functional impairments in vision are driven by retinal blood vessel leakage and macular edema^7^. More than 30% of patients with diabetes are reported to have DR^8^. Current treatments include laser photocoagulation, intravitreal steroids, and frequent intravitreal injections of vascular endothelial growth factor (VEGF) inhibitors, alongside blood sugar control. While effective, anti-VEGF therapies require repeated injections and can have off-target effects on retinal neuronal cells that express the signal-transducing VEGFR2^9–11^. Patients receiving systemic anti-VEGF treatments reported side effects, including hypertension, bleeding, vascular injury, proteinuria, impaired wound healing, and gastrointestinal perforation^12^. To circumvent these side-effects, anti-VEGF therapy for DR and AMD is administered intravitreally^13,14^.

Neovascular age-related macular degeneration (AMD) is another major retinal disease that leads to vision loss. Two distinct forms, dry and wet AMD are distinguished. Dry AMD, the more common form, is characterized by progressive thickening of Bruch’s membrane beneath the RPE, alongside gradual degeneration of photoreceptors, the RPE itself and the underlying choriocapillaris. This progressive damage to the surrounding tissue results in disruption of the BRB, contributing to retinal ischemia and inflammation. The more severe, wet form of AMD is driven by choroidal neovascularization (CNV), where abnormal blood vessels from the choroid invade the RPE and extend into the subretinal space, leading to fluid accumulation, increased vascular permeability, and vision loss. The breakdown of the BRB in wet AMD promotes the leakage of fluid and proteins from newly formed blood vessels, exacerbating retinal damage and accelerating disease progression.

The pathogenesis of AMD remains poorly understood and current treatment options are limited. In wet AMD, frontline therapeutics include intravitreal injections of VEGF inhibitors, such as bevacizumab and ranibizumab, which target pathological neovascularization and vascular leakage, are the mainstay of therapy^15^. More recently, bispecific antibodies targeting both VEGF and angiopoietin-2 (Ang-2) have been developed to improve treatment efficacy and reduce the frequency of injections^16^. Preclinical treatment with antibodies blocking the Slit2 receptors Roundabout1 and 2 reduces ocular inflammation and synergizes with anti-VEGF treatment in preventing mouse OIR and CNV^17^. However, despite these and other advances, AMD remains a major cause of irreversible blindness, highlighting the need for novel therapeutic strategies.

Netrin-1 is a key axon guidance molecule that has emerged as a crucial regulator of angiogenesis and BRB integrity^18,19^, and its dysregulation is implicated in DR pathogenesis^20–22^. Netrin-1 enhances retinal revascularization in OIR and diabetic models, while its knockdown using shRNA- reduced pathological angiogenesis^20,21^. Netrin-1 also modulates vascular permeability in DR^22^. MMP-9 cleaves Netrin-1 into a bioactive VI-V fragment that increases vascular permeability and diabetic macular edema, and inhibiting MMP-9 prevents this pathological permeability. In contrast, full length Netrin-1 preserves endothelial integrity^20,23^; thus underscoring the therapeutic potential of preserving Netrin-1 signaling in retinal vascular disease^22^.

We previously demonstrated that Netrin-1 maintains BRB integrity by activating endothelial Unc5b and enhancing Norrin–β-catenin signaling.^19^. Single cell RNA sequencing (scRNA-seq) of retinal ECs identified a BRB gene expression program that is regulated by Netrin-1 and Unc5b^19^. Additionally, UNC5B is essential for maintaining stability at endothelial cell–cell junctions in HUVECs and in retinal plexus branching^24^. These findings reveal the essential role of Netrin-1 in maintaining BRB integrity and suggest that targeting Netrin-1 signaling may restore barrier function and prevent vascular leakage, offering a novel therapeutic approach for DR and related neurovascular diseases.

Beyond the retina, Netrin-1 improves vascular integrity in the aged bone marrow, supporting hematopoietic recovery, while its loss increases vascular leak^25^. Netrin-1 is upregulated in cancers^26^ and its blockade using an anti-Netrin-1 antibody reduced human endometrial cancer tumor progression^27^. It has also shown to inhibit metastasis and improve survival in mouse pancreatic adenocarcinoma^28^. In diabetic models, Netrin-1 administration enhances insulin release from β-cells^29^. In atherosclerosis, Netrin-1 expression by lesional macrophages inhibits their emigration, contributing to plaque progression and chronic inflammation^30,31^. Blocking Netrin-1 signaling in these models promoted macrophage egress and reduced atherosclerotic burden, highlighting its dual role in both vascular stability and inflammatory retention. Together, these findings suggest that Netrin-1 plays a broader systemic role in vascular and tissue homeostasis, raising the possibility that modulating its signaling may offer therapeutic benefits across multiple disease contexts, including those affecting the retina.

In this work, we used genetic *NTN1* overexpression mice to determine effects on BRB integrity. We also employed the Oxygen Induced Retinopathy (OIR) and Laser-Induced CNV models of pathological leak, and we investigated the effect of *NTN1* overexpression on OIR induced vision impairment. We found that overexpressing Netrin1 stabilized the BRB in both physiological and pathological models of permeability, and improved vision recovery in OIR. In addition, we found that *NTN1* overexpression minimally and transiently affected retinal angiogenesis in an Unc5b independent manner, while its BRB stabilizing effects were Unc5b dependent. Together, these findings position Netrin-1-Unc5b signaling agonists as promising therapeutic candidates for restoring vascular integrity and visual function in retinal diseases characterized by BRB breakdown.

## MATERIALS AND METHODS

### Mouse models

All animal experiments were performed under a protocol approved by the Institutional Animal Care Use Committee of Yale University. All mice were of the C57BL/6 strain. Inducible Netrin-1 gain of function mice (gift from Patrick Mehlen, Lyon University) were generated by intercrossing loxStoplox;hNTN1^32^ with RosaCre^ERT2^ (Jackson laboratories) driver to induce ubiquitous overexpression of NTN1. Ntn1^fl/fl^ mice^33^ crossed with ROSACre^ERT2^ mice were described previously. Unc5b^fl/fl^ mice^34^ were bred with Cdh5Cre^ERT2^ (endothelial deletion) (Taconic, Biosciences), or with RosaCre^ERT2^ (global deletion).

### Western Blot

Retinas were dissected and frozen in liquid nitrogen and lysed in RIPA buffer (Research products, R26200-250.0) supplemented with protease and phosphatase inhibitor cocktails (Roche, 11836170001 and 4906845001) using a Tissue-Lyser (5 times 5min at 30 shakes/second). Protein lysates were centrifuged 15min at 13200rpm at 4°C and supernatants were isolated. Protein concentrations were quantified by BCA assay (Thermo Scientific, 23225) according to the manufacturer’s instructions. 30ug of protein was diluted in Laemmli buffer (Bio-Rad, 1610747) boiled at 95°C for 5min and loaded in 4-15% acrylamide gels (Bio-Rad, 5678084). After electrophoresis, proteins were transferred on a nitrocellulose membrane and incubated in TBS 0.1% Tween supplemented with 5% BSA for 1h to block non-specific binding. The following antibodies were incubated overnight at 4°C: Ntn1 (mouse anti-Netrin-1 R&D, AF1109, which shows cross-reactivity with recombinant human Netrin-1), β-actin (Sigma, A1978). Western blot densitometry analysis was conducted using ImageJ (version 1.54g).

### Tracer injection and immunostaining

Sulfo-NHS-biotin tracer was injected intraperitoneally in P12 mice at a concentration of 10mg/mL (300uL per mouse) and left to circulate for 1h.

Retinas from mice injected with tracers were dissected in 3.7% formaldehyde to ensure dye fixation within the tissue during retina dissection. Each retina was placed in 3.7% formaldehyde at room temperature for 10 minutes, stored in 100% methanol overnight, washed 3 x 10 minutes with PBS, dissected and incubated with specific antibodies in blocking buffer (1% fetal bovine serum, 3% BSA, 0.5% Triton X-100, 0.01% Na deoxycholate, 0.02% Sodium Azide in PBS at pH 7.4) overnight at 4°C. The following antibodies were used: Unc5b (Cell signaling, 13851S), Claudin-5-GFP (Invitrogen, 352588), Plvap (BD biosciences, 550563), Mfsd2a (Cell Signaling, 80302S), Esm1 (Biotechne, AF1999), Alexa-Fluor 647-Anti-ERG (abcam, AB196149), TER-119 (Invitrogen, 14-5921-82). IB4 as well as all corresponding secondary antibodies were purchased from Invitrogen as donkey anti-primary antibody coupled with either Alexa Fluor 488, 568 or 647. Streptavidin-Texas Red (Vector Laboratories, SA-5006-1) was used to detect sulfo-NHS-biotin. The next day, retinas were washed 3 x 10 min and incubated with secondary antibody for 2h at room temperature, then washed 3 x 10 min and post-fixed with 0.1% PFA for 10minutes and mounted using fluorescent mounting medium (DAKO, USA).

### Microscopy and image analysis

Confocal images were acquired on a laser scanning fluorescence microscope (Leica SP8) using the LASX software. 10X, 20X and 63X oil immersion objectives were used for acquisition using selective laser excitation (405, 488, 547, or 647 nm). Quantification of vascular outgrowth and quantification of pixel intensity were performed using the software ImageJ (version 1.54g) . Vascular density was quantified with the software Angiotool by quantifying the vascular surface area normalized to the total surface area. The superficial, intermediate and deep layers were separated and colored using ImageJ z-stack functions. Quantification of pixel intensity was performed using the software ImageJ. Avascular area in retinas from mice subjected to OIR was quantified on ImageJ by measuring the avascular area and normalizing to the total area of a retina. Claudin-5 was quantified using area of green pixels divided by area of all pixels. Plvap was quantified using area of magenta pixels divided by area of all pixels. Fluorescein leak was quantified by creating an area around a lesion based on a brightfield image and then quantifying intensity using the same ROI in the fluorescence channel. Quantification of TER-119 Integrated Density was measured on ImageJ. Integrated density is Mean intensity X area. CNV quantifications were done by creating an ROI of a lesion based on the brightfield image of the lesion, and then quantifying the fluorescence intensity of that ROI in the corresponding fluorescence channel.

### Oxygen-Induced Retinopathy

P7 mice were placed in a hyperoxia chamber with the dam. The chamber oxygen level was set to 75% and the chamber was closed from P7 to P12. On P12 the pups were returned to room air with a nursing foster mother. At P12, P13 and P14 mice were given TAM injections (200mg/mouse per day). Eyes were then assessed at P17 for the maximum neovascular response for OIR.

### Laser-induced choroidal neo-vascularization (CNV)

Adult mice were briefly anesthetized with ketamine (100mg/kg) and xylazine (10mg/kg). 1% Tropicamide (Somerset) was applied on eyes for 5 mins to dilate pupils. The tropicamide was then wiped off and a drop of GenTeal (Alcon Professional) was applied to the eyes. Four focal burns were administrated in each eye using a 532 nm argon ophthalmic laser via a slit lamp, being careful to avoid large blood vessels. One week post injury mice were anesthetized with ketamine (100mg/kg) and xylazine (10mg/kg) and intraperitoneally injected with 10mg of Sodium Fluorescein approximately 10 minutes prior to imaging, and lesions were imaged. Optimal coherence tomography (OCT) images were acquired using a Phoenix Micron IV. Shortly after, mice were sacrificed, enucleated and eyes were fixed in 4% PFA for 10 mins, before choroids were dissected for whole mount staining.

### Electroretinography

Electroretinography was performed as previously described^35^. Mice were dark-adapted overnight before being anesthetized with ketamine (100 mg/kg) and xylazine (10 mg/kg). To maintain body temperature, the mice were placed on a temperature regulating heating pad. Pupils were dilated by applying 1% Tropicamide to the eyes for 5 minutes, followed by positioning gold wire corneal electrodes (LKC Technologies, SKU: N30). Electroretinograms (ERGs) were recorded simultaneously from both eyes. Scotopic responses were obtained using single-flash stimuli ranging from ∼4.0 log cd·s/m² to +2.7 log cd·s/m². The a-wave amplitude was calculated as the negative deflection occurring within the first 60 milliseconds post-flash, while the b-wave amplitude was identified as the maximum positive deflection following the a-wave. A 55-Hz Bessel filter was applied to remove oscillatory potentials prior to measuring the b-wave amplitude. Mice were monitored for 24 hours after anesthesia administration.

## RESULTS

### NTN1 overexpression enhances BRB integrity during postnatal retinal development

Inducible Netrin-1 gain of function mice were generated by intercrossing mice harboring a human *NTN1* transgene (loxStoplox;hNTN1)^32^ with RosaCre^ERT2^ mice. Ubiquitous heterozygous *NTN1* overexpression was induced by TAM injections from P0-2 (hereafter *NTN1iGOF*) and TAM-injected Cre-negative NTN1fl/WT littermates were used as controls (**Fig.1A**). Subsequent qPCR analysis of P5 retinas with primers that detect human *NTN1* revealed increased *NTN1* mRNA levels in *NTN1iGOF* mice compared to littermate controls (**Fig. 1B**). Western blot of retina lysates using an antibody that detects mouse Netrin-1 and cross-reacts with human Netrin-1 confirmed increased protein levels in *NTN1iGOF* mice compared to littermate controls (**Fig.1C,D**).

**Figure 1.**
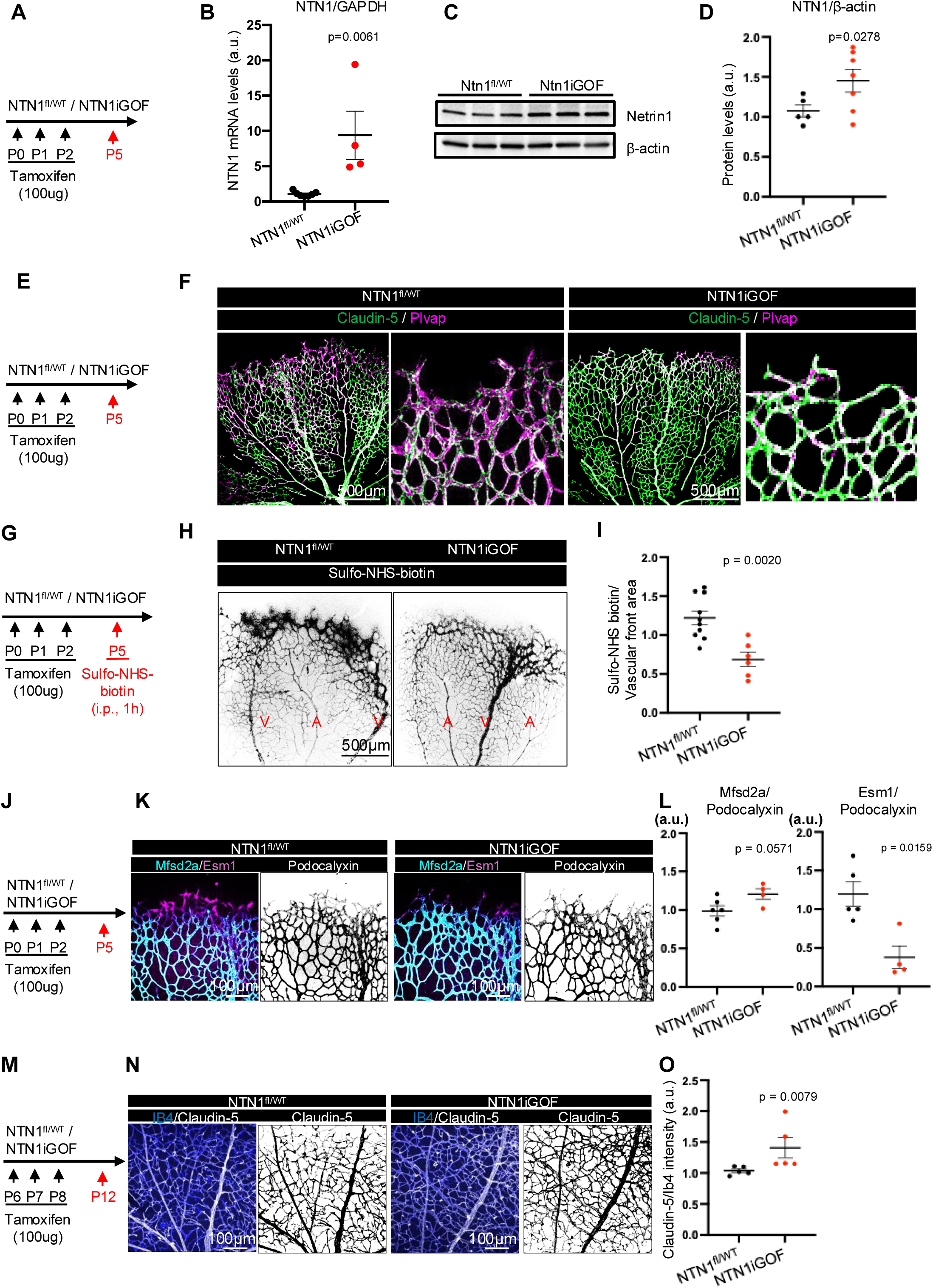
Netrin-1 overexpression prevents vascular leak in neonatal retinas (A) NTN1 overexpression strategy in neonates. (B) qPCR analysis of P5 retina mRNA extracts. (C) Western blot and quantification (D) of retina protein extracts 3 days post tamoxifen at P5. (E) NTN1 overexpression strategy in neonates. (F) immunofluorescence of whole-mount P5 retinas with the indicated antibodies. (G) Ntn1 overexpression and tracer injection strategy. (H) P5 retinas i.p. injected with Sulfo-NHS-biotin for 1h and (I) quantification of leak at the angiogenic front. (J) Deletion strategy (K) retina staining with indicated antibodies and (L) quantification of Mfsd2a-positive area relative to Pdxl-positive area and Esm1-positive area relative to Pdxl positive area. (M) Deletion strategy in P6 neonates. (N) P12 retinas stained with indicated antibodies and (O) quantification of Claudin-5 mean intensity/IB4 mean intensity. Data are shown as mean ± SEM. Two-sided Mann–Whitney U test was performed for statistical analysis.

As knockout of *Ntn1* and its receptor *Unc5b* destabilizes the BRB^19^, we anticipated that conversely, overexpressing *NTN1* would enhance barrier properties. Towards this goal, we investigated expression of Claudin-5, a tight junction protein, and Plvap, a permeability protein expressed in fenestrae and transcytotic vesicles in P5 retinas. The vascular front of *NTN1^fl/WT^* littermate control mice was Plvap+ and Claudin5- (**Fig. 1E, F**) and displayed higher Claudin-5 expression in more mature vessels closer to the optic nerve, and lower expression at the angiogenic front (**Fig. 1F**). Conversely Plvap expression was enriched in the leaky front but suppressed in mature vessels (**Fig. 1F**). By contrast, *NTN1iGOF* ECs at the vascular front converted to a Claudin-5+/Plvap- phenotype (**Fig. 1F**), consistent with acquisition of barrier function. Intraperitoneal (i.p.) injection of Sulfo-NHS biotin into P5 *NTN1iGOF* mice showed significantly reduced BRB leakage at the angiogenic front compared to Cre-negative littermate controls (**Fig. 1G,H,I**), attesting to enhanced vascular barrier formation in superficial tip cells. Probing for expression of the BRB-associated lipid transporter protein Mfsd2a revealed low levels of Mfsd2a in Esm1+ and Podocalyxin+ tip cells at the angiogenic front of Cre-negative littermate controls, as expected from prior studies^36,37^ (**Fig. 1J**). However, *NTN1iGOF* mice showed mildly increased levels of Mfsd2a in Podocalyxin+ tip ECs at the vascular front, and reduced Esm1 levels in Podocalyxin+ tip cells compared to controls, suggesting that molecules regulating barrier properties are induced^36^ (**Fig, 1K,L**). At P12, when retina vessels have acquired a functional BRB^19,37^, the *NTN1iGOF* mice still displayed increased Claudin-5 immunostaining levels compared to littermate controls (**Fig. 1M-O**) suggesting a reinforced endothelial barrier.

### Netrin-1 exerts a minimal and transient effect on retinal angiogenesis

Netrin-1 has been previously shown to affect angiogenesis^18,38–41^. Following TAM injections from P0-2, IB4 staining of P5 *NTN1iGOF* retinas revealed a small but significant increase in vascular outgrowth compared to littermate controls (**Fig. 2A,B)** but no change in vascular density (**Fig. 2C**). Conversely, *NTN1*iLOF (inducible loss-of-function/knockout) mice generated by intercrossing *NTN1*fl/fl mice with ROSA-Cre^ERT2^ and TAM-treated at P0-2 and analyzed at P5 like the GOF mice displayed no change in vascular outgrowth and a small but significant increase in vascular density compared to their littermate controls (**Fig. 2D-F**). We had previously reported that TAM treatment of *Ntn1iKO* mice between P6 and P12 produced no change in angiogenesis of the superficial, intermediate and deep vascular layers^19^. Likewise, P12 *NTN1iGOF* retinas from mice injected with TAM at P6 – P8 showed similar vascular density in superficial, intermediate and deep plexus compared to controls (**Fig. 2G-I**). Additionally, staining for the pericyte marker Pdgrfβ revealed no differences in pericyte coverage of IB4+ vessels between *NTN1iGOF* retinas and controls at P12 (**Fig. 2J,K)**. These data show that Netrin- 1 has a small and transient effect on early retinal angiogenesis, but sustained effects on BRB integrity.

**Figure 2.**
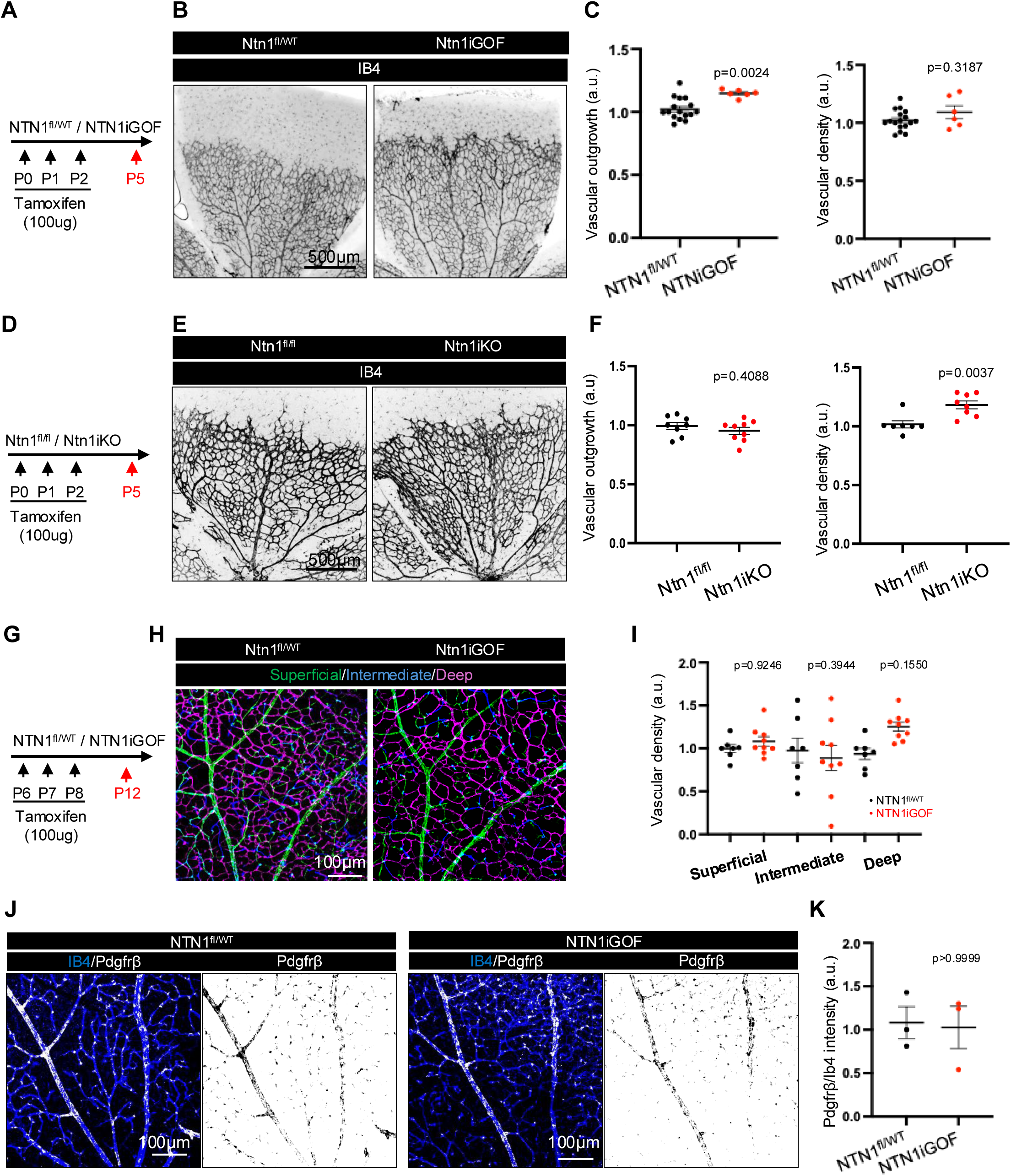
Netrin-1 promotes angiogenesis of the superficial retinal vascular plexus (A) NTN1 overexpression strategy in neonates. (B) Immunofluorescence of whole-mount P5 retinas stained with Ib4. (C) Quantification of vascular outgrowth and density. (D) Ntn1 gene deletion strategy in neonates. (E) Immunofluorescence of whole-mount P5 retinas stained with IB4. (F) Quantification of vascular outgrowth and density. (G) NTN1 overexpression strategy in P6 mice, (H) P12 retina staining and (I) quantification of IB4 labeled vascular layers in indicated genotypes. (J) Immunofluorescence of whole-mount P12 retinas stained with IB4 and Pdgfrβ and (K) quantification. Data are shown as mean ± SEM. Two-sided Mann-Whitney U test was performed for statistical analysis between two groups, ANOVA followed by Bonferroni’s multiple comparisons test was performed for statistical analysis between multiple groups.

### NTN1 overexpression rescues pathological vascular leakage in oxygen-induced retinopathy

To investigate whether *NTN1* overexpression could inhibit leak in ocular neovascular disease, we used the OIR model^42^ (**Fig. 3A**). Retinas of mice exposed to 75% oxygen (hyperoxia) between P7 to P12 develop vaso-obliteration and a capillary free avascular area in the center of the retina. Upon return to room air at P12, the relative hypoxia induced release of excessive amounts of angiogenic factors, which results in pathological sprouting and neovascular tuft formation, accompanied by leaky vasculature and hemorrhage. When *NTN1iGOF* was induced by TAM administration during the revascularization process, retinas displayed decreased avascular areas and decreased neovascular tuft areas compared to littermate controls (**Fig. 3A-C**). This indicates a shift toward productive angiogenesis, facilitating revascularization, while simultaneously suppressing pathological angiogenesis, which is characterized by disorganized, aberrant vessel growth in neovascular tufts (NVTs). To understand this process better, we focused on OIR tip cells that express Podocalyxin as well as high levels of Esm1 and low levels of Mfsd2a^36^. Podocalyxin+ tip cells in the *NTN1^fl/WT^* control OIR sprouting regions were positive for the tip-cell marker Esm1 with limited Mfsd2a expression, but Mfsd2a expression was significantly increased in *Ntn1iGOF* retinas, suggesting improved barrier stability (**Fig. 3D,E**). Additionally, while control littermate retinas demonstrated the presence of Plvap and diminished expression of Claudin-5 in the arterioles and venules of P17 OIR retinas, littermate *NTN1*iGOF retinas had increased Claudin-5 expression and suppressed Plvap expression (**Fig. 3F,G**). OIR retinas typically demonstrated leakage of Ter-119+ red blood cells as was seen in control *NTN^fl/WT^* retinas and quantified by measuring the mean intensity times area of Ter-119+ staining. NTN1iGOF mice displayed decreased Ter-119+ cells as compared to controls (**Fig. 3H,I**), demonstrating protection against OIR injury.

**Figure 3.**
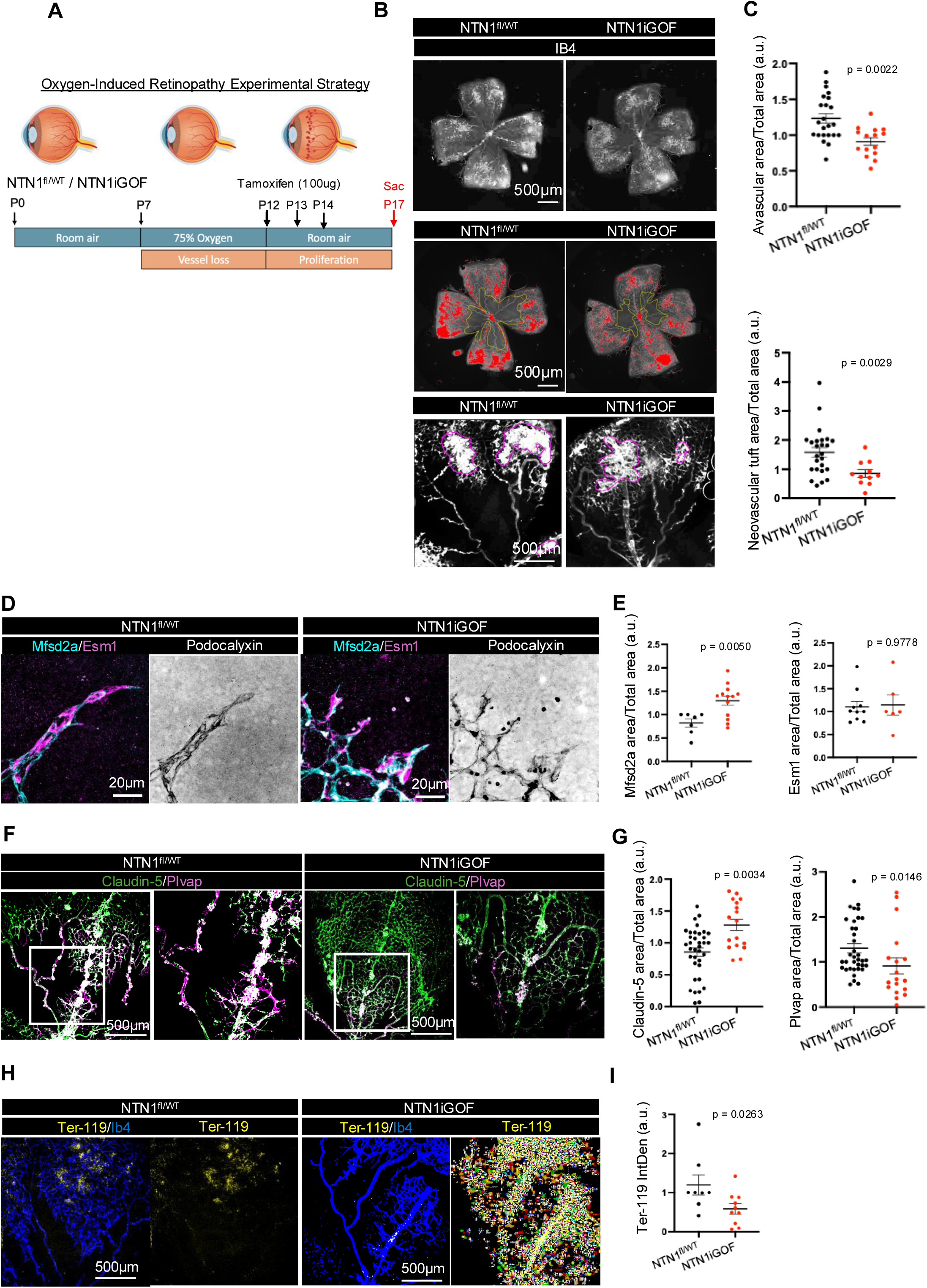
Oxygen-Induced Retinopathy in *NTN1iGOF* mice (A) OIR and gene deletion experimental strategy. (B) Whole-mount P17 retinas after OIR, one petal (top) and entire retina (bottom). Red indicates tufts and yellow boundary line indicates a vascular perimeter (C) Quantification of Avascular area and neovascular tufts. (D) High magnification images of P17 OIR sprouts stained with indicated antibodies and (E) quantification of Mfsd2a-positive area relative to Pdxl-positive area. (F) Whole-mount P17 retinas stained with indicated antibodies. (G) Quantification of immunostaining as indicated. (H) Whole-mount P17 retinas stained with indicated antibodies and (I) quantification of TER-119 Integrated Density (Mean intensity X area). Data are shown as mean ± SEM. Two-sided Mann–Whitney U test was performed for statistical analysis.

### Netrin-1 overexpression improves OIR-impaired vision

Retinal dysfunction and vision loss in the OIR retina are characterized by reduced electroretinogram (ERG) amplitudes, including photoreceptor (a-wave) and postsynaptic bipolar cell responses (b-wave)^42,43^. To determine whether a tightened barrier induced in *NTN1iGOF* mice improves retinal function in OIR mice, we conducted ERG recordings at P18 and P21. *NTN1iGOF* mice exhibited enhanced a- and b-wave amplitudes, with increasing light intensities, compared to littermate controls (**Fig. 4A-O**). Specifically, scotopic a-waves of *NTN1iGOF* showed significantly increased amplitudes compared to controls (**Fig 4.D,E**). Similarly, Scotopic b-waves showed a non-significant increase in step 4/5 (rod-only response) amplitude, and a significant increase in step 9/10 (mixed rod and cone response) amplitude (**Fig. 4.F,G,H**). Around P21 neovascularization in the retina spontaneously regresses and the retina vasculature recovers around P25^44^. When we measured ERG amplitudes in the same mice at P21 (**Fig. 4I**), we observed that responses of *NTN1iGOF* mice trended towards higher a and b wave amplitudes compared to littermate controls (**Fig. 4J-O**), but the values were not significantly different due to variability in the response (**Fig. 4L,N,O**). Overall, these data show that *NTN1iGOF* accelerates the recovery of visual acuity following OIR.

**Figure 4.**
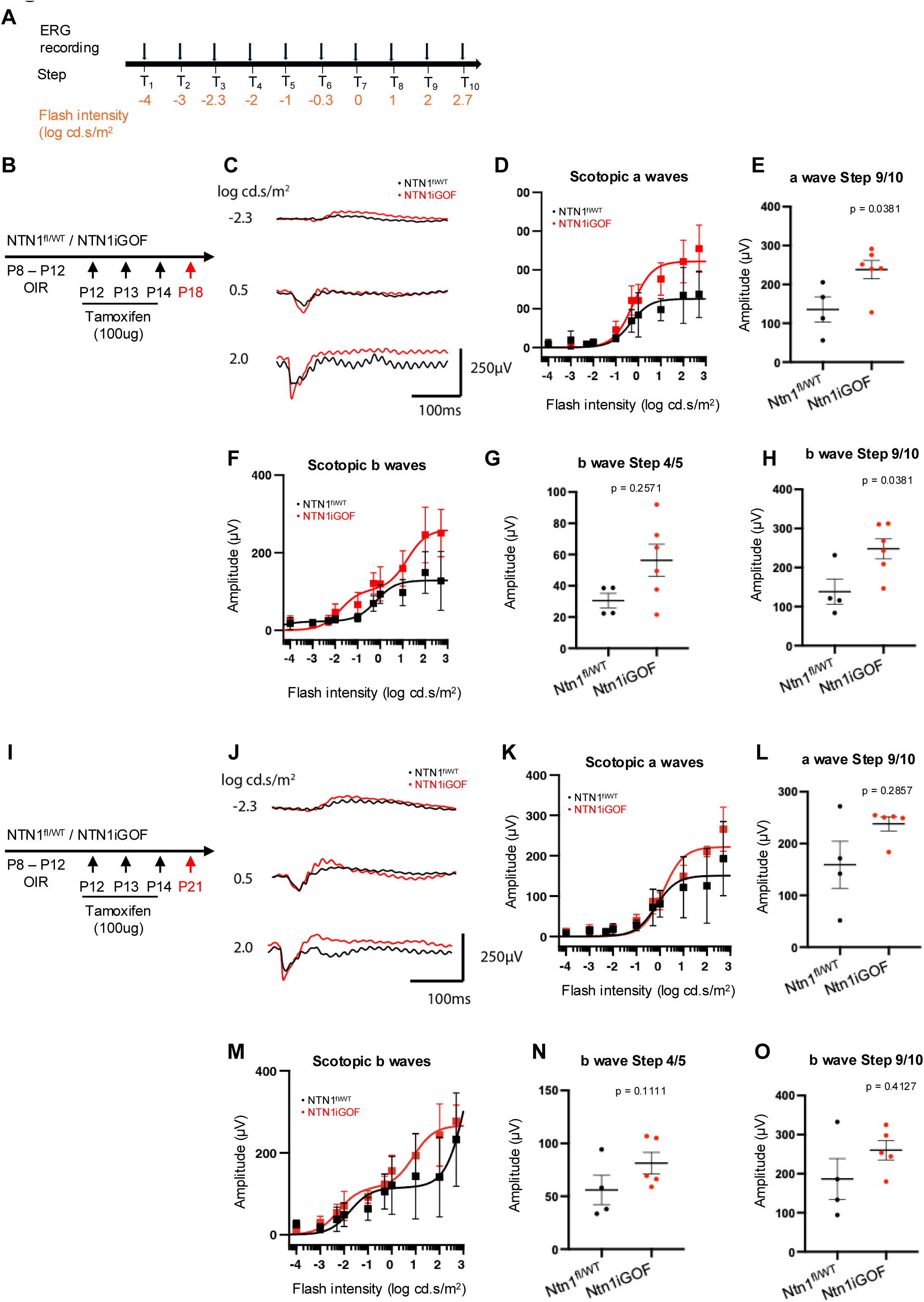
Netrin-1 overexpression improves OIR-impaired vision (A) ERG experimental outline. (B) NTN1 overexpression strategy and (C) Averaged scotopic combined traces at P18. (D) A wave amplitudes at P18 and (E) comparison of average response to step 9/10. (F) B wave amplitudes at P18 and (G,H) comparison at step 4/5 and step 9/10. (I) NTN1 overexpression strategy. (J) Averaged scotopic combined traces at P21. (K) A wave amplitudes at P18 and (L) comparison at step 9/10.(M) B wave amplitues at P18 and (N,O) comparison at step 4/5 and step 9/10. Data are shown as mean ± SEM. Two-sided Mann–Whitney U test was performed for statistical analysis.

### NTN1 overexpression reduces pathological leakage in choroidal neovascularization

To test effects of *NTN1* overexpression in another pathological model of leak, we used a laser-induced model that mimics neovascular/wet AMD^45^. The models consist of laser rupture of the Bruch’s membrane, which triggers the release of VEGF and results in the growth of new blood vessels from the choroid into the subretinal area. This neovascularization is accompanied by pathological leak assayed by measuring fluorescent tracer leak. Adult (P60) male *NTN1iGOF* mice and littermate controls were injected with TAM for 5 days and subjected to CNV three days later. CNV was induced by 4 distinct laser injuries per retina and vascular leak was assessed with fluorescein injection (i.p. 10min) 5 days later, which is reported to be the timepoint with maximal leak^46^ (**Fig. 5A,B**). Optical coherence tomography (OCT) imaging confirmed the CNV presence in both groups (**Fig 5B**). *NTN1*iGOF mice displayed markedly decreased fluorescence compared to littermate controls, as quantified by measuring mean fluorescein intensity per lesion and per eye (**Fig 5B,C**). Whole mount choroid staining showed no changes in IB4+ lesion size or smooth muscle cell coverage in *NTN1iGOF* mice compared to controls (**Fig. 5D,E**), indicating lack of an angiogenic effect in this model.

**Figure 5.**
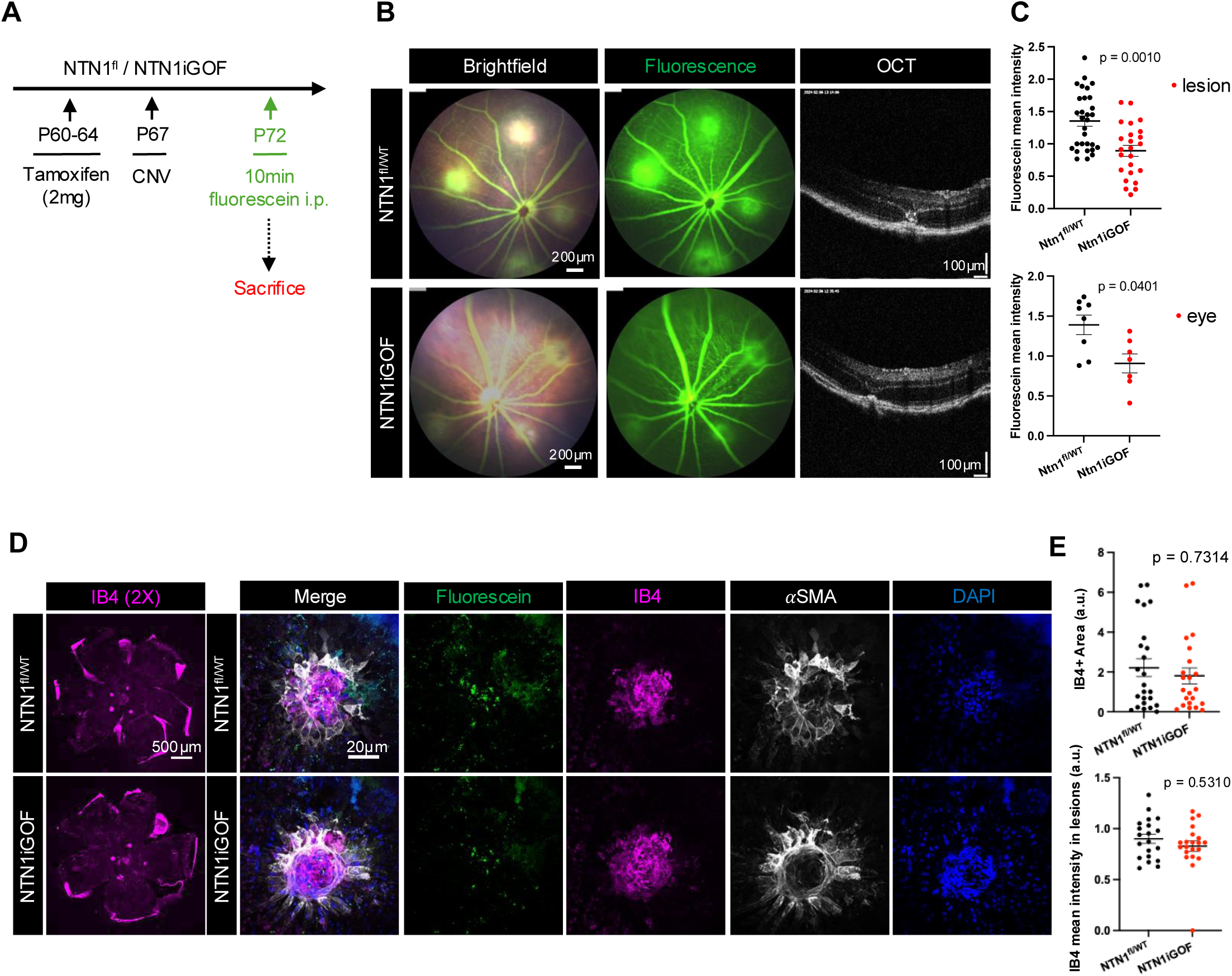
NTN1 overexpression rescues CNV-induced leak (A) NTN1 overexpression strategy and experimental design. (B) Fundus fluorescein angiography images 5-days post laser injury, and Optical Coherence Tomography (OCT) images 5-days post laser injury and (C) quantification of fluorescein mean intensity as indicated. (D) Whole mount staining of choroids with indicated antibodies and (E) Quantification. Data are shown as mean ± SEM. Two-sided Mann–Whitney U test was performed for statistical analysis.

### Unc5b Is Dispensable for Netrin-1–Mediated Angiogenesis but required for BRB integrity

To understand if Netrin-1 exerted its angiogenic effect on the superficial retinal vascular plexus via Unc5b, we queried single cell RNA sequencing data from P12 retinas for *Unc5b* expressing cell types. We observed Unc5b expression mainly in ECs, in mural cells, and in a few Muller glia cells^19^ (**Fig. 6A**). To determine whether Unc5b in pericytes or in Muller glia contributed to retinal angiogenesis or BRB function, we generated global Unc5b knockout mice (hereafter *Unc5biKO*) by intercrossing Unc5b^fl/fl^ and RosaCre^ERT2^.

**Fig 6.**
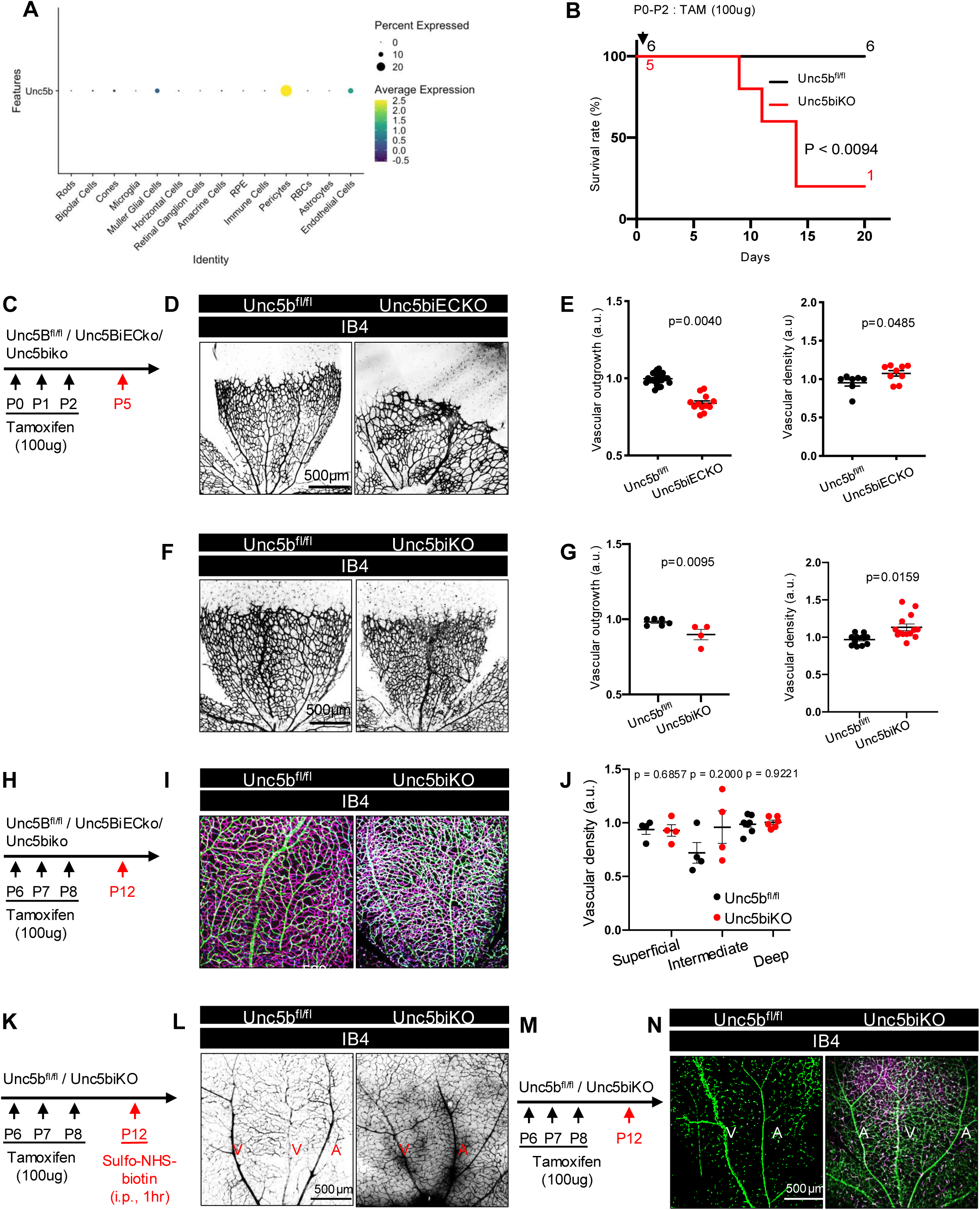
Effects of global and endothelial Unc5b deletion on retinal angiogenesis and the BRB (A) Unc5b gene expression in retinal scRNA-seq samples. (B) Survival curve after neonatal global Unc5b gene deletion. (C) Gene deletion strategy. (D) Whole-mount P5 retinas of indicated genotypes stained with Ib4 and (E) Quantification of vascular outgrowth and density. (F). Whole-mount P5 retinas of indicated genotypes stained with Ib4 and (G) Quantification of vascular outgrowth and density. (H) Gene deletion strategy. (I) IB4 staining and (J) quantification of superficial, intermediate, and deep layers at P12, in indicated genotypes. (K) Unc5b gene deletion and tracer injection strategy. (L). Whole-mount P12 retinas after i.p. injection with sulfo-NHS-biotin for 1h. (M) Unc5b gene deletion strategy. (N) Whole-mount P12 retinas stained with the indicated antibodies. Mantel-cox test was performed for survival curve statistical analysis in B. Data are shown as mean ± SEM. Two-sided Mann-Whitney U test was performed for statistical analysis between two groups, ANOVA followed by Bonferroni’s multiple comparisons test was performed for statistical analysis between multiple groups.

Neonatal TAM injections between P0-2 induced lethality of *Unc5biKO* mice by P15 (**Fig. 6B**) as previously reported with *Unc5biECKO* mice^34^. Immunostaining and western blot analysis of retinal lysates revealed efficient Unc5b deletion in *Unc5biKO* retinas (Supp **Fig. 1A-C**).

To assess retinal angiogenesis, we stained P5 retinal flatmounts from mice injected with TAM at P0-P2 with IB4 and measured vascular density and outgrowth. Endothelial Unc5b mutants displayed reduced vascular outgrowth and increased vascular density (**Fig. 6C-E**), as described previously in global Unc5b knockouts on an outbred background^41^. Global *Unc5b* deletions likewise reduced vascular outgrowth and increased vascular density (**Fig. 6C,F,G**). These phenotypes differ from *NTN1*iLOF mice (see Fig 2D-F) and lead us to conclude that Unc5b is not required for Netrin-1’s effect on angiogenesis. Global *Unc5b* deletion between P6-P8, and analysis of retinal vessel density at P12, revealed no significant differences between groups (**Fig.6H-J)**, highlighting that both Unc5b and NTN1 only affect angiogenesis of the superficial vascular plexus, but in a distinct manner.

To assess BRB permeability, we examined leakage of injected Sulfo-NHS-biotin in P12 mice after gene deletion between P6 and P8 (**Fig. 6K**). *Unc5biKO* mice exhibited BRB leakage (**Fig. 6M**), and ECs in the *Unc5b*iKO mice converted to a Claudin-5-/Plvap + low phenotype (**Fig. 6M-N**), demonstrating loss of BRB integrity.

## DISCUSSION

Herein, we report that TAM-inducible, global genetic *NTN1* overexpression in mice converted leaky vessels at the P5 angiogenic front into a non-leaky phenotype. Moreover, *NTN1* overexpression rescued pathological vessel leak in two ocular neovascular disease models, CNV and OIR. The barrier stabilizing effect in OIR improved electroretinogram recordings as seen by increased ERG amplitudes in P18 mice, attesting that *NTN1* promotes functional vision recovery along with BRB properties in this model.

At the molecular level, previous work revealed two distinct tip cell populations in the developing postnatal retina^36^. Superficial S tip cells are found at the leaky vascular front of P5 retinas, while D tip cells dive into the underlying neural retina to form two deeper vascular layers starting at P10. Superficial S tip cells are leaky to Sulfo-NHS biotin and express permeability proteins Esm1 and Plvap, while diving D tip cells are non-leaky and express Claudin5 and the lipid transporter Mfsd2a, which are two effectors of BRB integrity^36^. Neither Claudin5 nor Mfsd2a are expressed by superficial S tip cells, and neither Plvap nor Esm1 are expressed by diving D tip cells, hence expression of these proteins is mutually exclusive between leaky S-tip cells and BRB competent D-tip cells. The *NTN1iGOF* P5 superficial tip cells analyzed herein displayed loss of permeability proteins Plvap and Esm1 and upregulated Claudin5 and Mfsd2a, suggesting that *NTN1iGOF* converts leaky superficial tip cells into a phenotype that resembles BRB competent, non-leaky diving D-tip cells.

Pathological OIR tip cells express S-like mRNA profiles, including high Esm1 and low Mfsd2a expression^17,36^. *NTN1iGOF* upregulated expression of Mfsd2a and Claudin5 in OIR tip cells, while Plvap was suppressed compared to control OIR tips cells and Esm1 remained unchanged, suggesting that *NTN1iGOF* confers acquisition of some BRB properties. Furthermore, hemorrhagic injury was reduced, and visual recovery occurred earlier in *NTN1iGOF* mice when compared to controls.

In addition to effects on OIR vascular permeability, *NTN1iGOF* enhanced revascularization and decreased neovascular tuft formation. Netrin-1 as well as Unc5b and Neogenin expression levels were previously shown to increase in OIR retinas and pro-angiogenic effects of Netrin-1 in OIR were reported before ^47–49^,. One study reported that OIR-induced retinal neovascularization was successfully treated with Netrin-1 RNAi^48^ Another study conducted in mice lacking the related *Ntn4* gene showed that *Ntn4* homozygous global knockout mice subjected to OIR accelerated revascularization and recovery of visual acuity measured by ERG, with no effects of loss of *Ntn4* knockout on physiological retinal angiogenesis or in a CNV model were detected^50^. Here, *NTN1* loss or gain of function had weak and transient effects on angiogenesis. The most pronounced effect was seen in OIR angiogenesis, where avascular area and neovascular tuft formation were both reduced by *NTN1iGOF*, whereas neovascularization after CNV was unaffected. During development of the superficial vascular plexus between birth and P5, *NTN1iGOF* slightly but significantly increased vascular outgrowth but not vascular density, while *NTN1iLOF* slightly but significantly increased vascular density but not vascular outgrowth. Neither GOF (this work) nor LOF mutants^19^ displayed angiogenic defects when recombination was induced between P6 and P12, ie during formation of the deeper vascular layers of the retina. Thus, Netrin-1 has a small and transient effect on angiogenesis, while its effect on BRB integrity persists across all stages and models.

Our previous work showed that Netrin-1’s BRB stabilizing effects are mediated by endothelial Unc5b. Sc-RNA seq analysis showed that both *NTN1iLOF* and *Unc5biECKO* downregulated a common BRB gene expression program encoding tight junction proteins, nutrient and ion transporters, Wnt signaling effectors and others^19^. Herein, we extend these data by showing that global Unc5b mutants exhibit similar vascular phenotypes when compared to endothelial Unc5b mutants, including lethality and loss of BRB integrity. As both endothelial and global *Unc5b* increased vascular leak, additional Unc5b expression in retinal pericytes and Muller glia appears to play a minor role in BRB integrity when compared to endothelial Unc5b. Further studies using pericyte or Muller-glia specific Unc5b deletions should be conducted to confirm this finding.

By contrast to BRB integrity, we showed that NTN1 effects on angiogenesis did not require Unc5b, as phenotypes in the P5 retina of mutants were clearly distinct. *Unc5b* deletion in ECs or in all cells increased vascular density, confirming earlier work in global mutants and with function blocking antibodies^18,40,41^. Unc5b was identified as a Notch downstream effector and loss or gain of Unc5b limited the ability of Notch activation to regulate EC behaviors^24^. Endothelial-specific *Unc5b* deletion using the same lines employed here increased branching complexity in the developing retina^24^. While effects on vascular outgrowth was also increased in their series, for unclear reasons, both studies support a role for Unc5b in limiting vascular branching^24^. In addition to Netrin-1^40,51^, Unc5b binds Robo4^41,52^, Flrt2^53,54^ and Flrt3^53^ via its extracellular domain, and deletion of *Flrt3* phenocopied retinal hypervascularization observed in *Unc5biECko* retinas^55^. These data indicate that Unc5b mediated angiogenesis and barriergenesis are mediated by distinct ligands. Potential receptors mediating NTN1 effect on retinal angiogenesis include CD146^56^, which also binds VEGF^57^.

Dysregulation of Netrin-1 expression in human patients with DR was shown to contribute to vascular leak and macular edema^22^. In addition to current treatments for DR and AMD that rely on blocking VEGF-A activity^58–60^, recent studies developed agonists that induce BRB and BBB stabilization via activation of β-catenin^61,62^. Approaches to enhance Netrin-1 binding to endothelial Unc5b could synergize with such therapies and offer broad application through the regulation of both the BRB and the BBB.

## Statements & Declarations

### Author Contributions

Material preparation, data collection and analysis were performed by Jessica Furtado, Thomas Zapadka, Hyojin Park, Kevin Boyé and Jonathan Demb. First draft of the manuscript was written by Jessica Furtado and Anne Eichmann and all authors commented on previous versions of the manuscript. All authors read and approved the final manuscript.

### Data Availability

The data generated during and/or analysed during the current study can be made available on request.

### Ethics approval

All protocols and experimental procedures were approved by the Yale University Institutional Animal Care and Use Committee (IACUC).

### Consent to participate

No human subjects were involved in this study.

### Consent to publish

No human subjects were involved in this study.

---

**Supplementary Figure 1.**
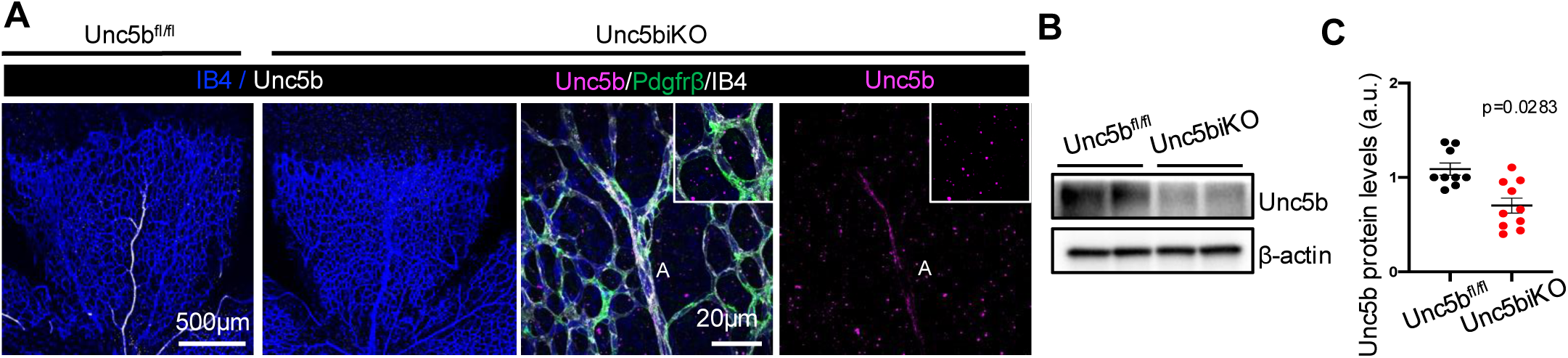
Validation of global Unc5b deletion (A) Whole-mount P5 retinas stained with indicated antibodies (B) Western blot of retina protein extracts probed for indicated proteins and blot quantification (C).

